# Attuning to song duels facilitates song-matching in nightingales

**DOI:** 10.1101/2025.04.12.648496

**Authors:** Giacomo Costalunga, Carolina Carpena Sanchez, Jonathan I. Benichov, Colin Cross, Jan Clemens, Daniela Vallentin

## Abstract

Complex vocal communication requires precise, flexible adjustments to vocal-motor commands for the production of coherent responses to auditory inputs. The mechanisms underlying how such sensorimotor integration controls vocal production is poorly understood. Here, we investigate the dynamics of territorial vocal duels and song-matching in common nightingales (*Luscinia megarhynchos*). Using a combination of field observations, playback experiments and accelerometry, we find that wild nightingales readily modified the acoustic features of their song, matched the rival’s song and engaged more vigorously against vocal opponents with a shared song repertoire. Furthermore, we demonstrate that neurons in the sensorimotor nucleus HVC are tuned to specific song stimuli, potentially enabling context-specific integration of auditory information to facilitate matched song production. These findings highlight the crucial role of song recognition in allowing nightingales to adjust their vocalizations in response to specific social situations, demonstrating the complexity and adaptability of their communication system. Moreover, this study provides a new framework and model system for investigating online sensorimotor processing during dynamic vocal interactions.

## Main Text

Behavioral flexibility can confer adaptive advantages by allowing the adjustment of motor outputs within variable environmental and social contexts. For example, human communication is characterized by remarkable diversity, whereby motor control of vocal organs, limbs, and facial muscles is modulated according to various contexts. Individuals readily adjust their expressions, body language and particularly speech patterns, tone, and content to suit different vocal partners and respond in real-time to their reactions (*1*–*4*). This inherent flexibility raises fundamental questions about how social context and communication partners drive the dynamic engagement in conversations, and how humans actively attune to certain interactions while dismissing others (*5*, *6*).

Acoustic interactions are widespread across other animals, with auditory-vocal integration driving both temporal and spectral coordination of vocalizations (*3*). Several species exhibit vocal turn-taking, precisely timing their utterances in response to others (*7–11*). One extraordinary example of vocal turn-taking is the production of cooperative song duets by some songbird species, in which vocal partners rapidly alternate, resulting in a unique shared song (*12–15*). While adjustments of the timing of vocalizations have been extensively investigated, evaluating changes in vocal content poses a greater challenge. This difficulty stems from the complexity of interpreting the quality of vocal responses in animal communication, where the "appropriateness" of a signal is often subjective and context-dependent (*16*).

This limitation can be overcome by studying species that engage in vocal matching (*17*–*20*). In these animals the matching vocal output can be used as a proxy for an appropriate, coherent response. A few songbird species perform song-matching, listening to and responding to their rivals by imitating their songs, primarily for territorial defense (*18*, *21*–*24*). In these duels, not only is the timing of the response crucial, but the vocal content must also be adjusted to produce an accurate match (*18, 23*). However, how the song content of the opponent influences song-matching behavior, as well as the underlying neural mechanisms facilitating this behavior, remains largely unexplored.

Pioneering neurophysiological studies in swamp sparrows (*Melospiza georgiana*), a species known for counter-singing using a limited song repertoire (2-5 songs), have revealed precise auditory-vocal mirroring of neural activity during song-matching (*25*, *26*). It was demonstrated that neurons in the premotor nucleus HVC (proper name), a key songbird brain region controlling the timing of song production (*27*, *28*) and implicated in song learning (*29*), exhibit the same activity patterns during both perception and production of single syllables. This "mirroring" of neural activity suggests a tight link between hearing and singing. However, it remains unknown how social context and rival song content influences song-matching behavior and how auditory information is integrated into song motor commands in birds with more complex vocal repertoires (*30*).

Here, we addressed the behavioral influence of rival song content and the neural auditory-motor integration processes for song-matching during song duels in Common nightingales (*Luscinia megarhynchos*). These songbirds have evoked a broad interest across centuries for their astonishing singing behavior. To defend territories and attract females, male nightingales use their extensive repertoires consisting of up to 270 different songs per bird to engage in complex song-matching duels, recursively countering each other (*23*, *31–33*).

We investigated how the song content of vocal opponents affects vocal content selection during song-matching behavior in nightingales in the wild. We conducted controlled playback experiments, which allowed us to precisely manipulate the social context by exposing wild birds to recordings of different conspecifics or themselves. When presented with their own songs, birds were more likely to counter-sing with matching vocal content, exhibited increased locomotor activity, and produced a more variable temporal song sequence. By contrast, nightingales engaged less with songs from conspecifics and more rarely matched their vocal content.

Furthermore, to explore the neural mechanisms underlying these rival-evoked behavioral shifts, we examined the activity of HVC neurons in response to song playbacks in hand-raised nightingales. This approach enabled us to shape their vocal repertoires and investigate how different song stimuli triggered changes in song motor pathway activity. We found that HVC neurons exhibited selective activity in response to songs within a bird’s own repertoire, suggesting a neural mechanism for recognizing and responding to specific songs, which could facilitate song-matching and vocal competition.

Our results demonstrate how social context affects flexible and complex vocal motor commands during vocal exchanges. Moreover, we introduce song matching behavior of nightingales as a novel framework to study real-time sensorimotor processes that directly influence vocal behaviors.

### Wild nightingales match rivals’ songs depending on the similarity of their song repertoires

To capture the natural dynamics of the nightingales’ complex counter-singing duels, we recorded nighttime vocal interactions between pairs of wild birds during their breeding season in Germany (Fig 1 A-B). Then, we performed controlled playback experiments to simulate song duels, exposing twenty male nightingales to songs of either an unfamiliar conspecific (conspecific’s songs playback, CON) or their own song sequence (bird’s own songs playback, BOS) (Fig 1 C).

**Fig. 1.**
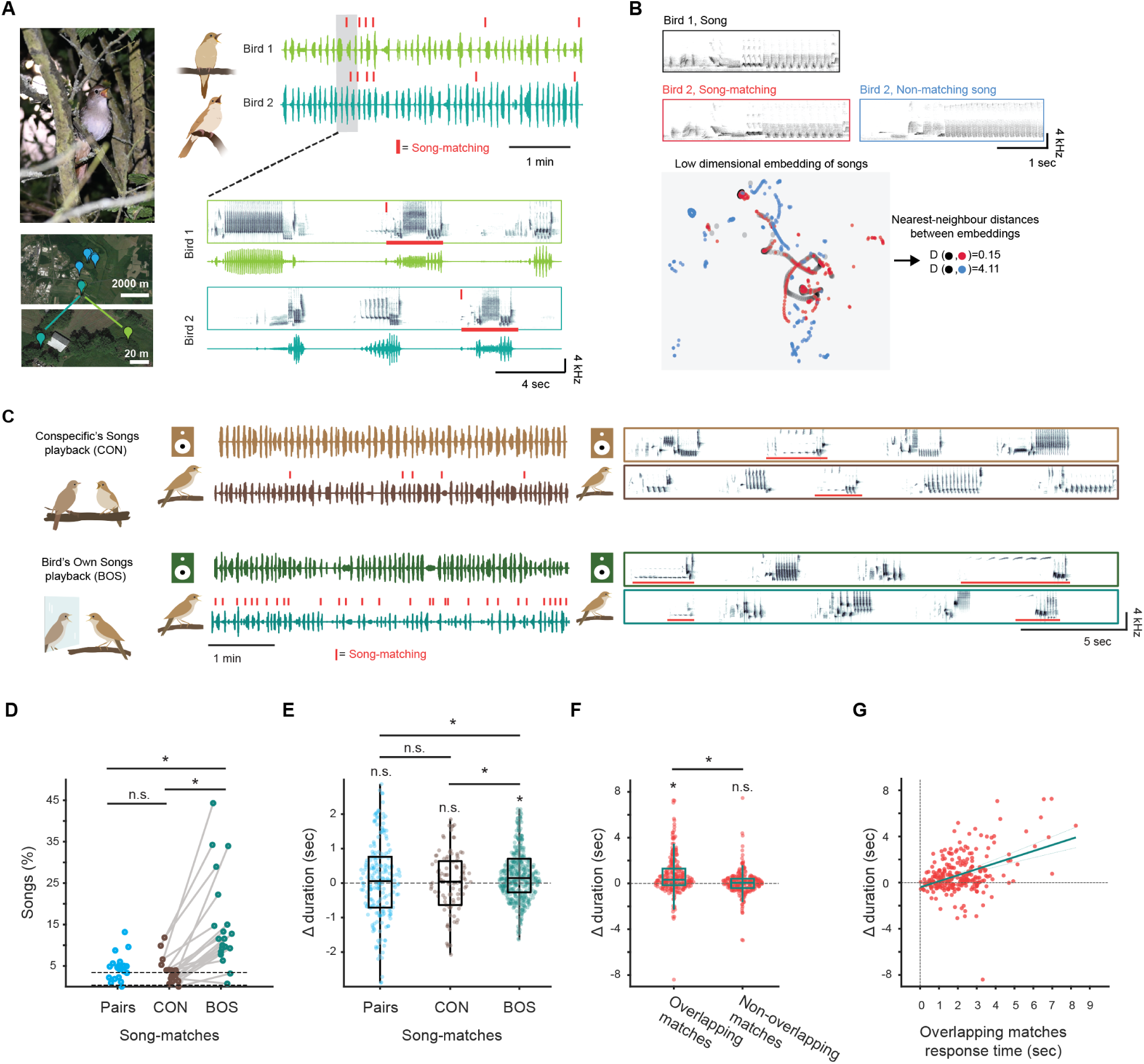
Wild nightingales match rivals’ songs depending on the similarity of the song repertoire. (**A**) Left: Singing nightingale; Location of example nightingale pairs in Germany (blue pins); Zoom-in on the territories of one example pair of nightingales (light and dark green pins). Right: Example of counter-singing duel between two nightingales in the wild. A closer analysis of the spectrograms and oscillograms of a vocal interaction highlighted in the dashed black box, shows that the last song of Bird 2 matches the second song of Bird 1. The matching songs are highlighted with a red line. See also Audio S1. (**B**) Spectrograms of a song of Bird 1 (black), and of matching (left, red) and non-matching songs (right, blue) from Bird 2. Spectrograms were embedded into a low dimensional space using UMAP, which transformed each song into a trajectory in this embedding space. Pairs of songs were considered matches if their trajectories’ nearest neighbor distance was small. (**C**) Left: Oscillograms of an excerpt from a playback experiment with one nightingale. During the playback stimulations, the nightingale’s songs were recorded simultaneously [top: conspecific (CON) playback, light brown, response below in dark brown; bottom: bird’s own song (BOS) playback, dark green, response in teal green below). The bird’s song-matches, in response to both playbacks, are highlighted with red tick marks. Right: example spectrograms of CON and BOS playbacks and bird responses, including song-matches highlighted with red lines. (**D**) Percentage of song-matching songs recorded: Pairs (light blue, 10 pairs, 20 birds), CON and BOS playback experiments (dark brown and teal green, respectively, 20 birds). Dashed line represents the chance level of random song-matching considering 100% repertoire sharing with the rival (neighboring male or playback). (Song-matching songs pairs= 4.6±3.1%, CON= 3.0±3.0%, BOS =11.5±11.5%, chance level of song matches= 4.0%).; Wilcoxon rank sum test, n.s. p=0.16, * p<0.001; Wilcoxon signed-rank test, p<0.001). (**E**) Distributions of Δ duration for song-matches during pair recordings and playback experiments. (Δ duration pairs= 0.0572±1.7047 sec, Δ duration CON matches= 0.0390±1.3234 sec; Δ duration BOS matches= 0.1438±1.4632 sec; Wilcoxon signed-rank test, n.s. p=0.5237, p=0.9608, * p<0.001; Wilcoxon rank sum test, n.s. p=0.6025, * p=0.0197, p=0.0205). Outliers in each distribution were removed to improve visualization. (**F**) Distributions of Δ duration for overlapping and non-overlapping song-matches to BOS playbacks (Δ duration overlapping matches BOS= 0.3220±1.6910 sec; non-overlapping matches BOS= 0.0533±1.0948sec; Wilcoxon rank sum test, * p<0.001, Wilcoxon signed-rank test, n.s. p=0.3584, * p<0.001). (**G**) Correlation between the Δ duration and the response latency of overlapping song-matches to BOS playbacks, aligned to matched song onset (Correlation: 261 overlapping matches BOS, Linear regression model, R-squared= 0.17, * p<0.001).

Previous playback experiments using a bird’s own song as a stimulus have produced varied results across songbird species ranging from neutral to aggressive responses (*21*, *34*, *35*). Owing to the competitive nature of nightingale counter-singing, we hypothesized that confronting a nightingale with its vocal *Doppelgänger* (BOS playbacks) would induce changes in singing behavior. We predicted that males would attend to the playbacks by increasing behavioral engagement and adjusting their vocalizations to compete with a rival that perfectly performed their own song.

To test this hypothesis, we analyzed the bird’s song matching behavior during vocal interactions. Therefore, we developed a semi-automated pipeline to detect matching songs by constructing continuous low-dimensional UMAP embeddings of the playbacks and responses from each bird (Fig 1B). While song-matching occurred equally often during naturalistic exchanges and CON playback stimuli, exposure to BOS playback triggered a fourfold increase (Fig 1 D), significantly surpassing what would be expected by chance with completely overlapping repertoires (see methods).

Song-matching has been proposed to be an aggressive signal in other species, particularly when responses temporally overlap with the stimulus (*36*). We found no consistent differences in response timing across playback conditions. Nightingales overlapped matches to CON and BOS playbacks with similar probabilities (BOS overlapping matches=51%; CON overlapping matches=42%) and did not precisely time their responses to the playbacks (range of response times aligned to stimulus onset, CON: 1.3 – 11.0 sec; BOS: 1.2 – 8.3 sec).

Different renditions of each song in a nightingale’s repertoire can be produced with variable repetitions of song syllables (Fig 1 B-C). Therefore, the birds produce the same song with different durations. To test whether the birds copy the same duration they just heard or a different one we correlated the duration of the matches to the duration of the stimuli. We found that BOS matches were consistently shorter than the stimuli whereas CON matches were similar in duration (Fig 1 E). Notably, overlapping matches were shorter than non-overlapping matches (Fig 1 F). Additionally, the longer the response latency of the overlapping responses to the onset of the matched song, the shorter the duration of the matching song (Fig 1 G).

Taken together, these findings suggest that the amount of song-matching is influenced by how strongly the opponent’s repertoire matches the content of the bird’s own songs. Additionally, the precise timing or song duration were less important than similarity in song content in counter-singing duels with a *Doppelgänger* rival.

### Repertoire similarity of rivals influences the temporal structure of nightingale song sequences

Beyond song-matching, we examined whether the social context influenced the timing of song production (Fig 2 A). Singing rate decreased (Fig. 2B, C) and became more variable (Fig. 2B, D) during BOS playbacks compared to CON playbacks, indicating that different dueling contexts affect the pacing of songs and silent gaps.

**Fig. 2.**
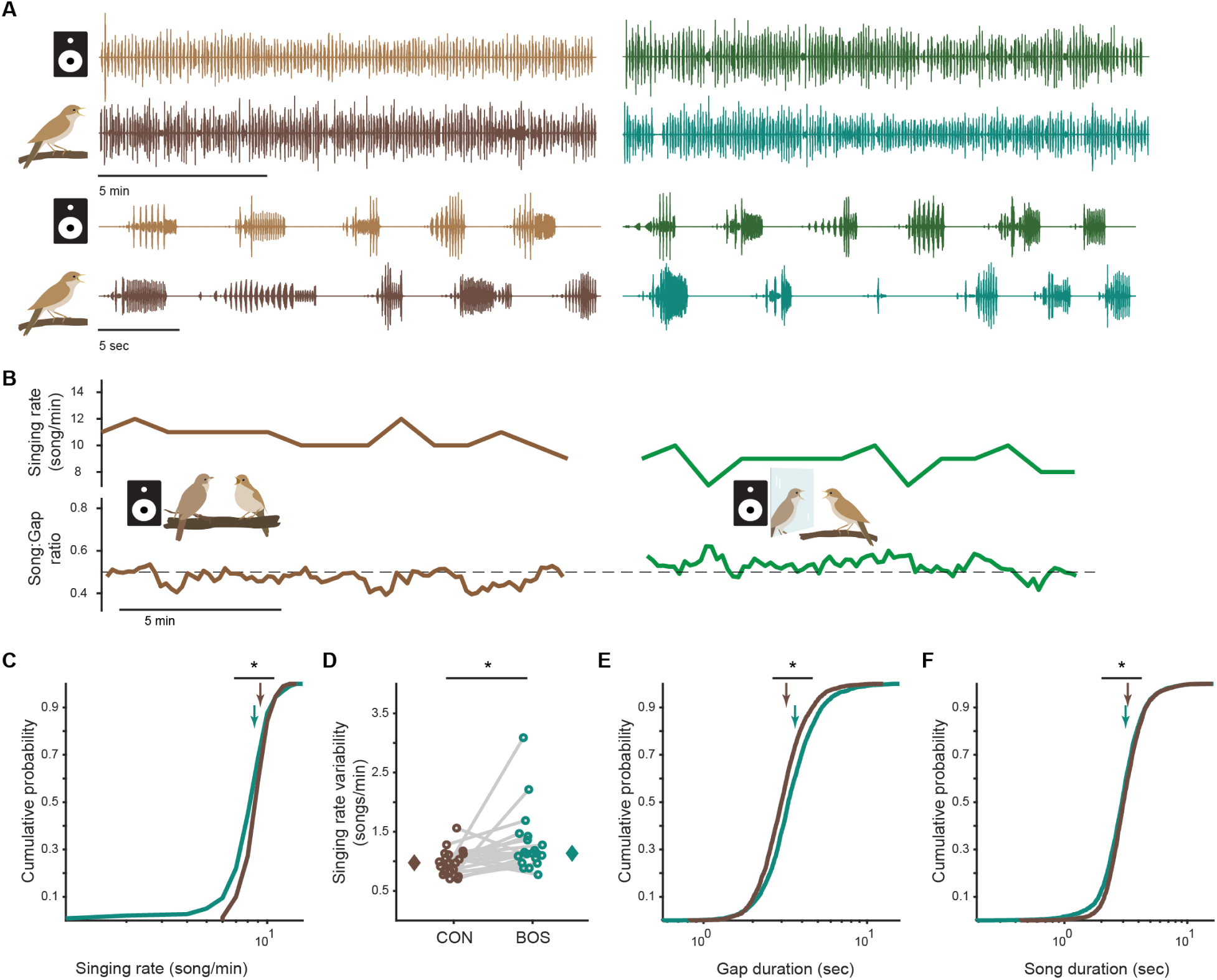
Temporal structure of nightingales’ song sequences during playbacks. (**A**) Top: Oscillograms from an example playback experiment with one nightingale (CON playback left, BOS playback right). Bottom: zoomed-in example snippet of the recording. (**B**) Top: Singing rate of an example nightingale during CON and BOS playbacks. Bottom: ratio between song and silent gap durations of the same bird. (**C**) Singing rate of 20 nightingales during CON and BOS playbacks. Arrows represent the mean values. (Mean singing rate CON= 9.2 songs/sec, BOS= 8.6 songs/sec; linear mixed-effect model, p<0.001). (**D**) Singing rate variability of nightingales singing against CON and BOS playbacks. Diamonds represent the median values. (Singing rate variability CON playback= 1.0±0.2 songs/sec, BOS playback= 1.1±0.5 songs/sec; Wilcoxon signed-rank test, * p= 0.019). (**E**) Duration of silent gaps of nightingales singing against CON and BOS playback. (Mean gap CON= 3.2 sec, BOS= 3.6 sec, linear mixed-effect model, * p<0.001). (**F**) Duration of songs of nightingales singing against CON and BOS playbacks. (Mean song duration CON= 3.3 sec, BOS= 3.1 sec, linear mixed-effect model, * p<0.001).

This shift in temporal dynamics was primarily driven by an increase in the duration of silent gaps between songs during BOS playbacks, compared to the shorter durations during CON playbacks (Fig 2 E). Moreover, in addition to shorter BOS matches as described above (Fig 1 E) also non matches were shorter during BOS playbacks compared to songs during CON playbacks (Fig 2 F). These finding indicate that song production dynamics are modulated by the similarity of the rivals’ songs to that of the bird’s own song repertoire.

Taken together, these results demonstrate that nightingales adjust multiple temporal parameters of their singing behavior depending on the repertoire content of the singing rivals. When countering their own songs, nightingales increased silent gap durations and shortened song durations despite increased song-matching. This suggests a strategy that prioritizes listening to the rival’s song before responding. This may facilitate more effective song-matching by allowing nightingales to accurately process and encode the preceding songs of their opponents.

### Singing duels involving the birds’ own song repertoire increase arousal

While studies have linked BOS playbacks to aggression in some songbird species, others have yielded mixed results depending on species and measurements utilized (*34*). To objectively assess the behavioral impact of BOS playbacks in nightingales, we quantified their overall movement during playback-induced singing duels. Since traditional visual observation methods are challenging for nocturnally active species living in dense vegetation like nightingales, we attached miniaturized accelerometers to individual birds to precisely track their movements in their natural habitats (Fig 3 A). This allowed us to quantify body acceleration and movement bouts beyond singing activity, providing a proxy for arousal and engagement during singing interactions with different playback stimuli (Fig 3 B-C).

**Fig. 3.**
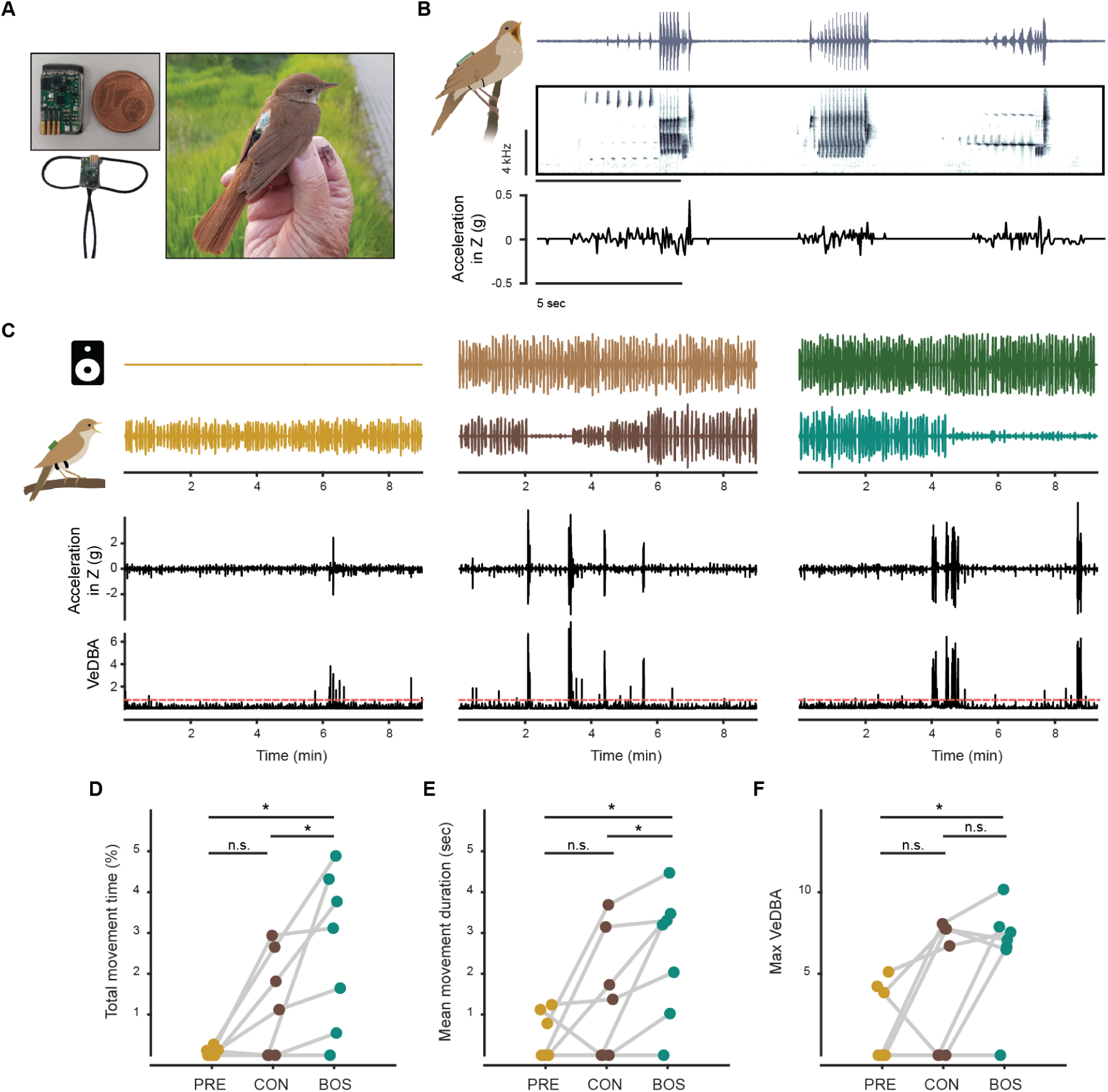
Accelerometry in the wild reveals behavioral arousal responses to song playbacks. (**A**) Lightweight accelerometer datalogger backpack to measure activity profile of wild nightingales. (**B**) Example of simultaneously recorded singing activity (top: oscillogram; middle: spectrogram) and acceleration profile (bottom) for three example songs of a wild nightingale. (**C**) Example of playback experiments aligned with accelerometer recordings of a nightingale. During the pre-playback phase (left), the recorded bird is not exposed to any song stimulus; During CON and BOS playbacks, movement bouts are visible in the accelerometer trace and VeDBA as high amplitude peaks, and correspond to changes in amplitude in the oscillogram of the bird’s singing activity resulting from changes in proximity to the directional microphone. Red dashed lines represent the threshold used to detect movements beyond singing from the VeDBA. (**D**) Total movement time as a percentage of each playback phase. (Total movement time: Pre-playback control= 0±0.1 %, CON playback= 1.1±1.3 %, BOS playback= 3.1±1.9 %; Wilcoxon signed-rank test, n.s. p= 0.13, * p= 0.031, n=7 birds). (**E**) Average duration of movement bouts during experimental playback phases. (Mean movement duration: Pre-playback control= 0±0.6 sec, CON playback= 1.4±1.5 sec, BOS playback= 3.2±1.6 sec; Wilcoxon signed-rank test, n.s. p= 0.19, * p= 0.031). (**F**) Maximum amplitude of acceleration in movement bouts during experimental playback phases. (Max acceleration: Pre-playback control= 0±2.4, CON playback= 6.7±4.1, BOS playback= 7.1±3.1; Wilcoxon signed-rank test, n.s. p= 0.31, p= 0.22, * p= 0.031).

During the pre-playback control phase, birds exhibited typical nocturnal singing behavior, remaining stationary on their preferred singing posts and singing continuously for extended periods (*31*) (Fig 3 C, left). Exposure to CON playbacks elicited occasional movement bouts in some birds (Fig 3 C, middle), likely reflecting attempts to locate the simulated rival. However, BOS playbacks triggered a drastic increase in movement bouts (Fig 3 C, right). Nightingales moved for more time in total, for longer duration bouts and faster (see methods) when counter-singing against BOS playbacks compared with CON playbacks and pre-playback controls (Fig 3 D - F).

These more vigorous responses to hearing playbacks of their own songs indicates heightened arousal and possibly a greater perceived threat from the *Doppelgänger* rival. This suggests a fine-tuned sensitivity to songs in their own repertoires, potentially serving as a key signal for behavioral engagement in song duels for territorial defense.

### HVC projection neurons in nightingales are tuned to different song stimuli

Our behavioral experiments with wild nightingales revealed that these birds differentially adjust the content and timing of their singing behavior in response to different dueling contexts, with song-matching behavior and arousal being most prevalent in response to *Doppelgänger* rivals. To understand the neural basis of this change in behavior, we investigated how auditory information is integrated into the song production pathway, focusing on the vocal-motor nucleus HVC. In songbirds, this nucleus generates the motor program for song production, exhibiting highly stereotyped activity that guides the temporal progression of singing (*27*, *28*, *37–39*). Moreover, it regulates precise response timing of birds engaging in call turn-taking (*7*, *10*, *40*). Although it receives direct auditory inputs (*41*, *42*) and shows auditory-evoked activity in response to song playbacks in juvenile zebra finches (*29*, *43*, *44*), these auditory evoked responses are suppressed in adult birds that do not perform song-matching (*26*, *29*, *45*, *46*). This raises the question of whether HVC activity may be specifically modulated in nightingales and other song-matching species to enable their complex song interactions (*25*).

We bred nightingales in controlled laboratory settings and hand-raised nightingale chicks, thus controlling for songs in their adult repertoire (Fig 4 A). Once these birds reached adulthood, we performed intra- and extracellular electrophysiological recordings from HVC projection neurons in awake listening male nightingales (37 HVC projection neurons, 4 birds) while either exposing them to playbacks of novel songs from wild conspecific (CON playbacks) or stimuli from their own song repertoire (BOS playbacks), approximating the playback paradigm used in the field experiments with wild birds (Fig 5 B-C). We observed that some neurons exhibit action potentials at specific time-points throughout the playback presentation, and that this evoked activity was not related to single syllables in the song stimuli (Fig 4 B-C).

**Fig. 4.**
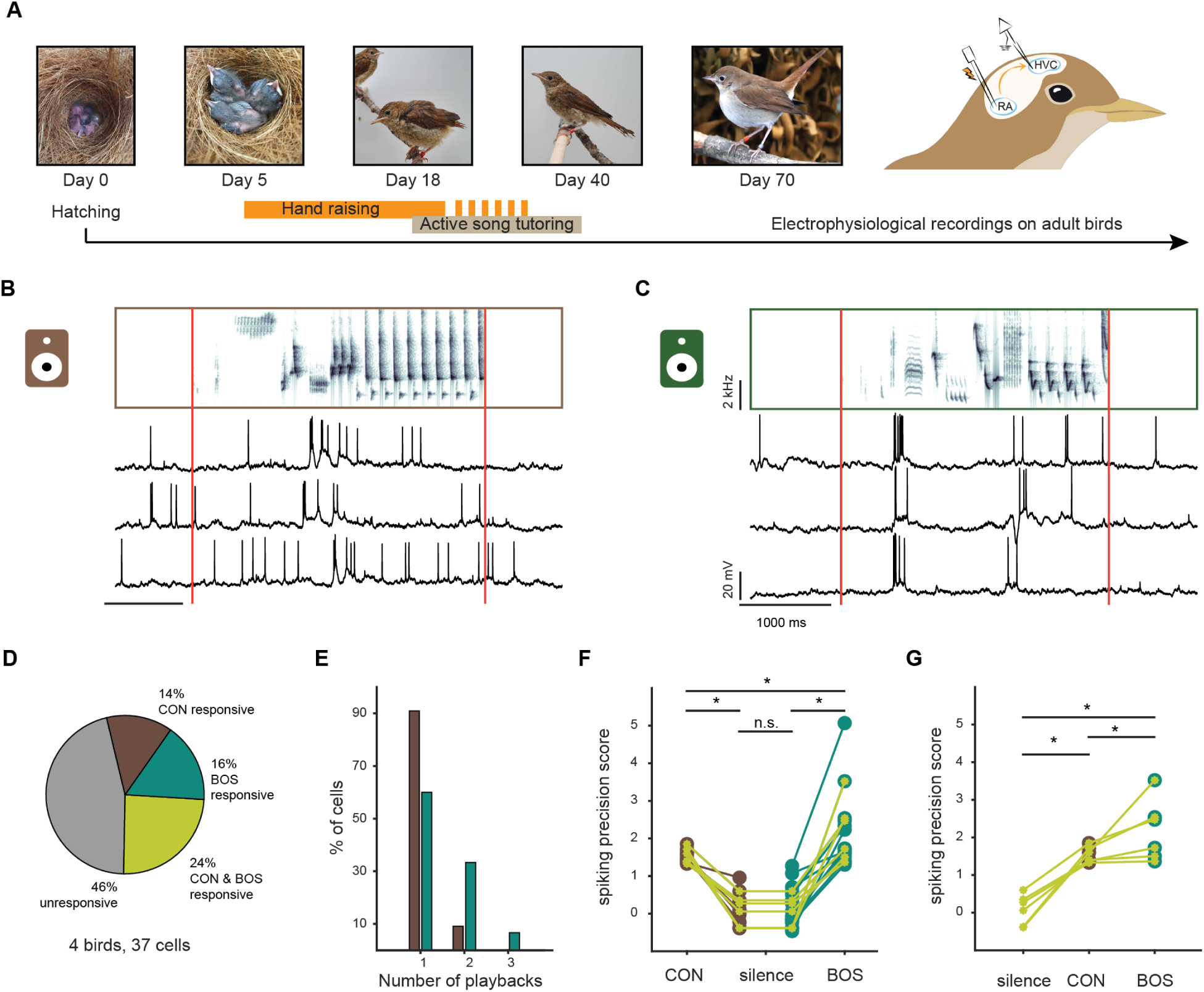
Electrophysiological recordings of HVC neurons in hand-raised nightingales reveal song stimuli tuning. (**A**) Timeline for electrophysiological experiments in hand raised nightingales. (**B**) Example recording of an HVC projection neuron during CON playback. Top: Spectrogram of song playback. Bottom: Example traces of the membrane potential of the neuron during three renditions of the playback. Red vertical lines indicate playback onsets and offsets. (**C**) As in (B), for the same HVC projection neuron during BOS playback. (**D**) Proportion of recorded neurons that are responsive to the different types of playbacks. (**E**) Number of different song playbacks eliciting responses in the stimuli-tuned neurons. (**F**) Spiking precision of the responsive cells, during silence and the different playback types. For cells responsive to more than one playback of the same type, the spiking precision score in response to the song playback eliciting the most precise activity is indicated (Precision score BOS playback= 1.7±1.0, CON playback= 1.5±0.1, silence= 0.1±0.4; Wilcoxon signed-rank test, * silence-CON p<0.001, * silence-BOS p<0.001, * CON-BOS p= 0.033, n.s. silence p= 1). (**G**) Spiking precisions of the cells responsive to both CON and BOS playbacks (Precision score BOS playback= 2.1±0.8, CON playback= 1.5±0.2, silence= 0.2±0.4; Wilcoxon signed-rank test, * p= 0.031).

Next, we quantified spike timing precision of HVC neurons, as an indication of stereotyped firing activity, time-locked to the song playback presentation. As a baseline, we computed the spiking precision during periods of silence preceding the playbacks. Then, we compared this baseline to the spiking precision observed during the presentation of the different playbacks. We found that a subset of the recorded neurons showed precise spiking in response to song playbacks, with some selectively tuned to CON, BOS, or both stimuli (Fig 4 D). Sometimes neurons responded to more than one song stimulus (Fig 4 E). This demonstrates that auditory exposure alone can drive stereotyped activity within the HVC of awake adult nightingales (Fig 4 B-C) and highlights the persistent auditory processing capabilities of this premotor nucleus. To quantify the effect of the different playbacks on spike timing precision, we compared the responses to CON and BOS stimuli. BOS playbacks elicited more precise time-locked activity in HVC projection neurons, both at the population level (Fig 4 F) and within those individual neurons responsive to both song types (Fig 4 G). This higher degree of stereotyped auditory-evoked activity in response to a bird’s own songs suggests direct processing of these behaviorally salient stimuli, facilitating song-matching.

## Discussion

We showed that a rival’s shared song repertoire influences singing dynamics and behavior in nightingales. Exposure to *Doppelgänger* playbacks (BOS) increased song-matching, led to variable temporal song dynamics and heightened general arousal. This contrasts with less engaged responses to unfamiliar songs from conspecifics, highlighting the existence of a song recognition system that goes beyond simple species identification. Our results demonstrate context-dependent vocal adaptability in a songbird species that uses its vocalizations to dynamically song-match different singing opponents.

Our findings contribute to the ongoing debate about the function and perception of vocalizations in songbirds (*34*). While previous behavioral studies in birds such as song sparrows showed mixed responses to BOS playbacks, our results in nightingales, with their extensive song repertoires and complex territorial dynamics, suggest a strong behavioral salience of these stimuli.

In humans, listening to an individuals’ own voice can elicit the phenomenon of “voice confrontation”, provoking general discomfort and aversion (*47*) and similar reactions can arise from visually depicted facsimiles of a person that fall into the “uncanny-valley” - a concept describing the dip in comfort levels when observing human-like entities that appear almost, but not quite, real, often evoking feelings of unease or revulsion (*48*). The “voice confrontation” phenomenon is partially explained by the difference in hearing sounds from external sources or those that are self-generated, resulting in a mismatch in auditory self-perception. Whether this extends to other animal species remains unknown, and the standard behavioral tests for self-perception in animals including birds rely on responses to visual representation of individuals in mirror self-recognition tests (*34*, *49*). In this study, we did not aim to test for self-perception or voice confrontation in nightingales. Notably, the birds did not seem to recognize themselves in the BOS playbacks, but rather consider BOS playbacks as relevant competitors and engaged with them with enhanced responses.

For nightingales, being confronted with a virtual *Doppelgänger* playback seems to represent an extremely behaviorally salient form of territorial dispute. Here, the exaggerated rival stimulus, in the form of BOS playback, leads to exaggerated behavioral responses in a manner that is perhaps analogous to responses to “supernormal stimuli” (*50*). A supernormal stimulus is an exaggerated artificial stimulus that elicits a stronger response from an animal than the natural stimulus for which the response evolved. These stimuli have been investigated behaviorally in several animal species, mostly for visual stimuli, including male butterflies trying to mate with more colorful dummies compared to real females and fish fighting wooden models that are more highly colored than real conspecifics. Here, BOS playbacks could represent “supernormal” auditory stimuli for counter-singing nightingales, driving increased territorial responses compared to CON playbacks.

The observed precise tuning of HVC neurons to BOS playbacks provides neurophysiological support for this heightened sensitivity to own vocalizations. The stereotyped activity that we observed in HVC neurons of hand-reared nightingales, which are typically suppressed in adult zebra finches (*29*, *45*), suggests a specialized mechanism for encoding and responding to songs within an individual’s own repertoire. This builds on previous work on counter-singing swamp sparrows which have a small song repertoire consisting of 2-5 different songs whereby each song is comprised of one single trilled, multi-note syllable (*26*, *51*). In these birds a subset of HVC neurons responded to a single song by exhibiting precise action potentials locked to each repletion of the syllable (*25*, *26*). In contrast, the song repertoire of a nightingale contains up to 280 different songs whereby each individual song consists of up to 12 different syllables (*31*). In these birds, we observed that HVC neurons responded to multiple songs and not to single syllables within each song, suggesting a role of this vocal-motor nucleus for distinguishing entire songs in one’s own repertoire from unfamiliar song content. Moreover, in swamp sparrows the same HVC neurons responding to the playback of a syllable discharged action potentials at exactly the same timepoints when the birds were singing the same song syllable (*26*). In light of this, our observations that song matching behavior is increased when nightingales are exposed to their own vocal repertoire leads us to speculate that the entire song matching process in nightingales is facilitated by priming HVC neurons to generate entire songs in real time.

By demonstrating complex, context-dependent vocal behavior and revealing its neural correlates, this study establishes the nightingale as a valuable model for studying the neuroethology of vocal communication. Song-matching offers a unique opportunity to dissect the neural mechanisms of distinct sensory and motor phases of sensorimotor integration (*9*), while the clear behavioral output allows for unbiased assessment of spectral adjustments and vocal adaptation during interactive communication. Future research can leverage this model to explore the neural mechanisms underlying vocal learning, social recognition, and strategic vocal communication in unprecedented detail.

## Acknowledgments

We would like to thank A. Proß, N. Mysuru, M. Pexirra, L. Bistere, M. Sensi and N. Hein for their help with fieldwork; Susanne Seltmann for helping with animal permits and logistics; H. Hultsch, and D. Todt for useful discussions at early stages of the project; T. Eliav for providing helpful comments to the early version of the manuscript; all the nightingale breeders for support with the laboratory birds; and A. Costalunga for the help with graphical design.

## Funding

HORIZON EUROPE European Research Council (ERC)-2017-StG-757459 MIDNIGHT (DV)

Deutsche Forschungsgemeinschaft VA742/2-1 (DV)

Deutsche Forschungsgemeinschaft 327654276–SFB 1315 (DV)

Joachim Herz Stiftung Add-on Fellowships for Interdisciplinary Life Science (GC) Deutsche Forschungsgemeinschaft BE7545/1-1 (JIB)

HORIZON EUROPE European Research Council (ERC) StG 851210 NeuSoSen (JC)

Deutsche Forschungsgemeinschaft SPP2205 596/2-2 430158535 (JC)

## Author contributions

Conceptualization: GC, DV

Methodology: GC, CCS, CC, JC, DV

Investigation: GC, CCS, JIB, JC, DV

Visualization: GC, CCS, JC, DV

Funding acquisition: GC, JIB, JC, DV

Project administration: GC, DV

Supervision: DV

Writing – original draft: GC, DV

Writing – review & editing: GC, CCS, JIB, JC, DV

## Materials and Methods

### Animal model

For experiments with wild birds during breeding season, we studied a wild population of Common nightingales (*Luscinia megarhynchos*) in semi-urban areas of Teltow-Fläming in the southwestern part of Brandenburg, Germany during mating season (April - May) between 2020 and 2023. All experiments in the wild were performed in accordance with the local authorities (Landesamt für Umwelt - Land Brandenburg LFU-N4 4730/14+5#181132/2021; Landesamt für Arbeitsschutz, Verbraucherschutz und Gesundheit - Land Brandenburg AZ2347-3-2021). For experiment with nightingales in laboratory settings, we used nightingales obtained from private breeders and descending from captive birds (See below). All Laboratory experiments were performed with the ethical approval of the Max Planck Institute for Biological Intelligence and the Regierungspräsidium Oberbayern (ROB-55.2-2532.Vet_02-18-182; ROB-55.2-2532.Vet_02-21-201).

### Vocal recordings

Audio recordings (16-bit precision at 44.1 kHz sampling rate) were made from wild male nightingales during the mating seasons of 2020, 2021, 2022 and 2023. Nocturnal recording sessions were conducted between 12pm and 4am CET+1, when nightingales performed counter-singing. Each nightingale was recorded with a directional parabolic microphone equipped with a windshield (Stereo MK3, Telinga, Sweden), connected to a battery-driven pre-amplifier (Roland Duo-Capture EX, Roland, Japan) and a laptop computer. Signals were acquired using Audacity v.2.4.2 (https://www.audacityteam.org/) and saved as stereo.wav files.

### Paired recordings

Both nightingales within a pair of interacting birds (10 pairs, 2 from 2020 and 8 from 2021, in total 20 birds) were recorded simultaneously. Microphones were placed ∼5-15 meters away from the birds singing from two locations within their territory, spaced ∼30-80 meters apart.

### Song playback experiment

We presented song playbacks to 20 wild nightingales in 2021. For the experiments, a speaker (JBL, Harman International Industries, USA) was connected to the pre-amplifier and placed at ∼10 meters from the singing bird and playbacks were broadcasted at a sound pressure level of ∼80 dB (A) (dB re. 20 μPa measured with a sound pressure level meter SPL (SL-100, Voltcraft, Germany) to simulate a bird singing at ∼95 dB (A) and located ∼6 meters apart.

Each nightingale was recorded during one session. A period of 14.4456 - 38.6084min (mean 23.7927±7.4262min) was recorded while the bird was singing without playback stimulation (pre-playback control). The playback phase consisted of a natural sequence of 152 songs recorded in 2020 from a wild male (∼15 minutes, Conspecific playback, CON) and of a natural sequence of 111-247 songs (16.3331±3.406 minutes) recorded during the pre-control phase of the session (Bird’s Own Songs playback, BOS). All birds were exposed with the CON playback first, followed by a silence time of ∼15 minutes before presentation of the BOS playback.

### Accelerometer experiments

To monitor activity in free-ranging wild nightingales exposed to playbacks, we equipped 17 males (7 in 2022 and 10 in 2023) with miniaturized accelerometer dataloggers (Axy5 XS, Technosmart, Italy), recording 3D acceleration 24/7 for several days at 25 Hz with a range of 8g. Birds were caught with the use of mist nets in proximity of their territories, in the early morning or in the late afternoon, using luring nightingale tapes. After assessment of the health status of the birds, morphological measures were collected and the accelerometers were secured on the birds. In addition, birds were ringed with unique color bands for individual recognition. The lightweight device (∼1.5 g) was fixed as a backpack on the back of the bird with rubber-band straps around both femurs and the abdomen. Birds were released in their territories, and post hoc analysis of the actograms revealed fast adjustment to normal activity in all birds. To allow the birds to recover from catching and adapt to the tags, playback experiments with birds carrying an accelerometer were performed at least 24 hours after first capture, if birds were singing at night. The loggers were retrieved few days later, by re-catching the animals in their territories. We successfully recapture and retrieved data from 14 birds, out of which 7 birds were successfully presented with both CON and BOS playbacks and thus included in the study.

### Housing, breeding and song tutoring of captive nightingales

Animals obtained from breeders were singly housed in aviaries partially communicating with the outdoors, and were provided with food and water *ad libitum*. During breeding season, pairs of males and females were formed by removing separating walls of the aviaries. Five days post hatching (dph), nightingales’ chicks were removed from the nest and transferred to indoor sound-attenuated chambers together with their siblings. The chicks were hand raised for approximately 20 days, until they were able to feed autonomously. Around 18 dph, we actively tutored the male chicks with selected nightingale songs, following established methods. Preceding surgical procedure, adult nightingales were placed in sound-attenuated boxes and their songs were recorded.

### Neural recordings

#### Surgery

Nightingales were anesthetized with isoflurane (1–3% in oxygen). After incision of the skin, the skull was exposed and the trabecular bone structure above the area of interest was removed with a dental drill (carbide bur, FG ¼, Johnson-Promident). The centers of the robust nucleus of the arcopallium (RA) and HVC, were located based on stereotactic coordinates and small craniotomies and durotomies were performed at the cranial surface targets. A chlorided silver ground wire (0.001”, California Fine Wires) was implanted above the cerebellum as ground. For nucleus targeting, a carbon fiber microelectrode (Kation Scientific) was advanced to the coordinates of the nucleus RA to verify its location via detection of characteristic high frequency tonic multiunit activity. For antidromic identification of HVC, a bipolar stimulating electrode was implanted into RA and antidromic responses were confirmed via the targeting electrode in HVC. A custom-made stainless-steel head plate was affixed to the skull using dental acrylic (Paladur, Heraeus). The craniotomies were protected until experiments were conducted using a silicone elastomer (Kwik-Cast; WPI). Animals were returned to their home cage 24 hours post-surgery and were monitored to ensure full recovery before experiments commenced.

#### Playback stimuli presentation

For measuring neural responses to songs, we presented playbacks at ∼65 dB through a speaker placed in front of the head-fixed birds. The presented stimuli were recordings of conspecific nightingale songs (CON playbacks) or songs of the bird’s own repertoire (BOS playbacks). For intracellular recordings, audio files were manually triggered after a successful stable intracellular recording was achieved. For extracellular recordings, audio files consisted of stereo tracks. One of the stereo channels was played via the speakers, the second channel consisted of a 10 ms 15 kHz pulse that was simultaneously played at the onset of each stimulus to ensure subsequent uniform alignment of playbacks with the neural data. This 15 kHz pulse was not directed to the speakers, but only used to trigger a TTL pulse for data alignment (see below).

#### Intracellular recordings

Recordings were performed after allowing for one recovery day following surgery. Awake birds were held in a soft foam restraint and head-fixed on the implanted head plate. Sharp glass pipette electrodes (borosilicate glass with filament, 0.1 mm diameter, impedance: 70-150MΩ) were prepared with a horizontal micropipette puller (P-97; Sutter Instruments) and backfilled with 3 M potassium acetate. The craniotomy over HVC was exposed, protected with 10% PBS and the brain nucleus was targeted based on stereotaxic coordinates using a micromanipulator (model MP-285A, Sutter Instrument). In the vicinity of HVC, the pipette was advanced in ∼5μm steps until positioned against a somatic membrane, as indicated by an increase in electrode resistance. A brief (∼10 ms) “buzz” pulse was then delivered to penetrate the cell membrane. Once a stable recording (spike height: >40 mV, resting membrane potential: <-50 mV) was achieved, song playbacks were presented. Membrane potential signals were amplified (Neuro Data IR183A, Cygnus Technology, Inc.), digitized at 40 kHz using a National Instruments acquisition interface, and recorded with custom MATLAB software. Based on the firing rate and the spike waveform of the recorded neurons, we concluded that none of them were inhibitory interneurons.

#### Extracellular recordings

A linear 16-channel silicon probe (NeuroNexus) was lowered into HVC using a micromanipulator (Sutter Instruments). Neural activity was digitized at a sampling rate of 30 kHz on an Intan RHD2132 headstage and acquired with an RHD Recording Controller (Intan Technologies). A TTL pulse was triggered by the 15 kHz tone at the onset of each playback presentation using an Arduino Uno, and delivered to the RHD Recording Controller for acquisition alongside the neural data.

### Analysis and statistics

All statistical tests used are specified in the main text. Values are reported as mean ± std, if not noted otherwise.

#### Audio data

Audio recordings from Germany were processed and analyzed using Audacity, Avisoft SASLab Pro 5.2 (R. Specht, Berlin, Germany) and Matlab.

For pair recordings, 16-33 min (28.14±5.87 min) of continuous recordings without contaminating background sounds were selected for data analysis.

For all recordings, stereo files were divided into mono files and a high pass-filtered (frequency=1000 Hz, roll-off=6 dB, High-Pass Filter build-in function of Audacity) and noise-reduced to remove environmental noises (noise reduction=12dB, sensitivity=6.00, frequency smoothing=3, noise reduction build-in function of Audacity).

Avisoft was used for semi-automatic segmentation of songs, using the built-in functions Create section labels from waveform events with parameters adjusted case-by-case. Onset and offset of detected songs were visually inspected and manually corrected, if needed.

#### Song-matching detection

Songs were matched using a segmentation-free and label-free approach by embedding each song’s spectrogram into a 4-dimensional UMAP space (*52, 53*) and comparing songs based on nearest-neighbor distances in this embedding space.

The power spectrograms with a logarithmic mel frequency scale were computed over 10 ms windows with a temporal resolution of 1 ms. All frequency bands below 0.63 and above 16.5 kHz were discarded as they were outside the range of most nightingale songs and therefore only contained environmental noise. We then subtracted the background noise, estimated as the median power over time for each frequency band, and log-transformed the spectrogram values. Spectrograms were then smoothed in time and frequency using a Gaussian kernel with a standard deviation of 5 ms and 3 mel frequency bands, and all values below 0 were set to 0. Finally, the temporal resolution was further reduced by a factor of 10, leading to spectrograms with a temporal resolution of 10 ms.

We then delay-embedded each spectrogram using a history of 40 spectrogram frames (400 ms) and reduced the dimensionality of the space from 40 to 4 dimensions using UMAP (*54*). For each individual bird, we learned an embedding using the playback stimuli for that bird. The bird’s playbacks and responses were then embedded into that learned UMAP space, resulting in each song’s spectrogram being represented as a 4D trajectory.

Pairs of songs were matched based on their nearest-neighbor distance in the embedding space. For each song pair, we computed the pairwise Euclidean distances between the embedded frames. For each frame in the stimulus, we then identified the distance of the nearest response frame in embedding space. We took the median of these frame-wise nearest-neighbor distances as a distance measure between song pairs. Songs closer than 0.7 were considered matches.

Song pairs with a small nearest-neighbor distance in this embedding space were considered matches. This approach produced reliable song matching (median F1 score over birds 80%). Song-matching performance was then further improved by manual proofreading.

#### Accelerometer data

The leg-loop harness used to secure the accelerometer resulted in positioning the device on the bird’s lower back. Considered the small size of nightingales, we expected to record acceleration from approximately the center of mass of the birds. Accelerometer data was recorded for the 3D axis (X, Y, Z). We calculated Vectorial Dynamic Body Acceleration (VeDBA) from the accelerometer data, a standard measure of overall movement activity calculated as the square root of the summed vector measures from an accelerometer. The data corresponding to the different phases of the playback experiment was extracted by aligning audio recordings with the acceleration profiles. The data was demeaned and a threshold was applied to capture movement that exceeded the singing activity. The thresholding was validated using the audio recordings, to include all movements that did not depend on the singing activity of the birds.

#### Neural data

To evaluate the temporal precision of neuronal firing, a precision score was calculated following established methods (*29*). In brief, the timing differences between all spike pairs across trials of the same auditory stimulus were computed, and the average of these differences was used as the precision score. To assess statistical significance, a permutation test was performed by randomly shuffling spikes across trials 1,000 times while maintaining the overall timing distribution. The mean latency difference from each shuffled dataset was then calculated to create a null distribution. Cells were classified as responsive if their original precision score fell outside the central 95% of this shuffled distribution.

